# Disentangling the diet composition of Arctic shorebirds’ chicks provides a new perspective on trophic mismatches

**DOI:** 10.1101/2022.10.10.511540

**Authors:** Mikhail K. Zhemchuzhnikov, Elena A. Zhemchuzhnikova, Thomas K. Lameris, Judith van Bleijswijk, Job ten Horn, Mikhail Soloviev, Viktor Golovnyuk, Maria Sukhova, Anastasiya Popovkina, Dmitry Kutcherov, Jan A. van Gils

## Abstract

With rapid climatic changes over the past decades, organisms living in seasonal environments are suggested to increasingly face trophic mismatches: the disruption of synchrony between different trophic levels due to a different phenological response to increasing temperatures. Strong effects of mismatches are especially expected in the Arctic region, where climatic changes are most rapid. Nevertheless, relatively few studies have found strong evidence for trophic mismatches between the breeding period of Arctic-breeding shorebirds and the arthropod prey on which they rely. Here we argue that this is potentially caused by a generalisation of trophic interactions. While many studies have measured the mismatch relative to the peak in abundance of all available arthropod species, we use metabarcoding of prey items in faeces to show that chicks of four different shorebird species (red knot, curlew sandpiper, little stint, and red phalarope) strongly differ in their arthropod diet. Three out of the four species feed on arthropods peaking in availability five-ten days before the overall arthropod peak which had implications for the calculations of trophic mismatches. We conclude that ignoring diet selectivity hampers our understanding of phenological mismatches.

## Introduction

With a warming climate, animals are considered to increasingly face trophic mismatches due to the growing phenological asynchrony between trophic levels (Thackeray et al. 2010). Organisms at higher trophic levels are shown to advance slower than those at lower trophic levels (Thackeray et al. 2016), which may lead to the weakening of the interaction strength between them. As such, mismatches are expected to result in fitness reductions, for example reductions in growth and survival for offspring growing up after the main seasonal food peak (Reed, Jenouvrier, and Visser 2013). However, there is no clear link between degree of mismatch that a population experiences and its fitness reductions (Zhemchuzhnikov et al. 2021), and some studies found no effects on fitness at all (e.g. Reneerkens et al. 2016).

Aside from the various factors due to which mismatches may have (or not have) fitness consequences, it is crucial to measure trophic interactions reliably. The structure of complex trophic webs, e.g., those which include a generalist predator and several potential prey types, heavily depends on the predator dietary preference (Mallord et al. 2016). However, when the exact prey preference is unknown, all potential food items are usually summed up to a single overall measure of abundance (e.g. Reneerkens et al. 2016). Such oversimplification of trophic interaction may lead to wrong conclusions, e.g., when estimating the degree of phenological synchrony between the trophic levels.

The same kind of generalization is often applied to the chicks of Arctic-breeding shorebirds and their arthropod prey (Schmidt et al. 2017; Reneerkens et al. 2016). Growth and survival of chicks are considered to depend on the biomass of all available arthropod species. Chicks growing up after the overall biomass peak generally experience reduced growth rates (Lameris et al. 2022) and survival chances (Meyer et al. 2021), yet other studies do not find fitness reductions for late-hatching chicks (Corkery, Nol, and Mckinnon 2019; Reneerkens et al. 2016). Before we can conclude that not all populations are equally sensitive to mismatches, it is essential to know which part of the total arthropod biomass is relevant to a certain bird species, as chicks of different shorebird species may not all rely on the same type of prey (Baker 1977; Gerik 2018; Holmes and Pitelka 1968).

Relatively few studies are dedicated to the analysis of diet in Arctic shorebirds and their chicks (Drury 1961; Wirta et al. 2015; Baker 1977; Holmes and Pitelka 1968; Gerik 2018; Holmes 1966). Most of them have been performed using microscopic analyses of the prey remains in the excrement and the digestive system (Baker 1977; Holmes and Pitelka 1968; Holmes 1966; Drury 1961). Modern molecular genetic tools, such as metabarcoding, can be successfully applied for detailed diet analyses of insectivorous birds (Wirta et al. 2016; Gerik 2018), including accurate estimates of not only the presence but also the relative abundance of different species in the diet (Rytkönen et al. 2019; Verkuil et al. 2022). Here we use a metabarcoding approach to determine the diet composition of chicks of four shorebird species with wide distributions in the Russian Arctic, including red knot *Calidris canutus*, curlew sandpiper *Calidris ferruginea*, red phalarope *Phalaropus fulicarius* and little stint *Calidris minuta*. We use these data to describe the (mis)match between phenology of these wader species (a) with the phenology of the overall arthropod abundance and (b) with the phenology of their key prey items.

## Methods

### Study area

The study was conducted near Knipovich Bay (76°04’ N, 98°32’ E), on the Taimyr Peninsula in the central Russian Arctic. The study area can be defined as Arctic tundra with alternating valleys and hills in the prostrate dwarf-shrub subzone (Walker et al. 2005).

### Arthropod sampling and biomass estimation

In 2018 and 2019, arthropod abundance was measured from late June till late July, during the chick rearing period of shorebirds, by using yellow round pitfall traps (⍰ = 9 cm), following Reneerkens et al. (2016). Traps were filled with propylene-glycol mixed with water in 1:1 ratio. We sampled arthropods in two (in 2018) or three (in 2019) grids, 0.64 km^2^ each. Each grid consisted of ten traps: nine of them were located in nodes of the grid at intervals of 400 m, and one was randomly allocated inside the grid between two neighbouring nodes. We emptied traps every five days and stored the containment in ethanol. This resulted in 120 (in 2018) and 180 (2019) samples in total. Arthropods in each sample were identified up to a family level in the lab. We did not collect Collembola and Acari, as their contribution to biomass is low, although in some low wet places there was a significant visible representation of these groups. From each sample we measured the length of a random representative for each family. This was used to estimate biomass based on the length-weight relationships in different arthropod groups, following Hodar (1996) and Sample et al. (1993).

### Collecting phenological data on shorebird nests

In both years active nest searching was conducted during the egg-laying and incubation phase from early until late June. The methods included observation of birds during egg-laying or after being flushed from the nest during incubation, rope dragging, and tracking birds to their nests with the aid of radio transmitters deployed before incubation (the latter only for red knots). Upon finding a nest, the incubation stage was determined using flotation tests. We tracked the fate of nests of shorebirds by revisiting them at least once, a few days before expected hatching. If the nest was not predated, we revisited it again on the expected hatching date to determine its ultimate fate. Hatch date was determined as either expected hatch date based on flotation tests (for predated, deserted nests and nests with unknown fate) or observed hatch date. Thus, the hatch date was established for 205 (100 in 2018, 105 in 2019) nests of little stint, 55 (14, 41) nests of red phalarope, 40 (18, 22) nests of curlew sandpiper and 16 (10, 6) nests of red knot.

### Shorebird broods and faeces sampling

Chicks were captured from late June till late July. Shorebird broods were either detected visually (curlew sandpipers, red phalaropes and little stints), or found by attracting males with playback of chick calls (red knots). To collect faeces, we placed each chick into a separate section of a thermo-insulated bag (curlew sandpipers, red phalaropes and little stints) or placed them together without separation (red knots). After a maximum of 15 minutes, chicks were released near their parent. Droppings found in the bag after release were stored in ethanol and transported to the lab for molecular-genetic analysis. In total 153 dropping samples were collected and analysed, among which 52 of broods (52 broods) of red knot, 23 (14 broods) of curlew sandpiper, 15 (8 broods) of red phalarope and 63 (33 broods) of little stint. Red knot faeces were sampled both in 2018 (22 samples) and in 2019 (30 samples), while all the other shorebird faeces were sampled in 2019 only. Sampling dates were distributed equally across the whole sampling period in both years.

### Barcoding method description

Our protocol followed the methods described in Verkuil et al. (2022) with respect to DNA extraction, PCR protocol with primers on the CO-I gene and settings for the bioinformatics workflow based on OTU clustering with a 97% identity cut-off. Taxonomy assignment was done based on a custom database containing 69 newly derived Sanger sequences of all insect morphotypes caught in traps during the fieldwork campaign in Taimyr plus 1337 sequences of Arctic insects taken from GenBank. A detailed description of the molecular genetic methods and the pipeline for processing the data as well as the reference database are given in Supplementary files (S1-3). As the result we obtained the data on the diet of shorebird chicks on a brood level. The arthropods assignments were grouped at family level (except for the Araneae, Collembola and Acari).

### Statistical analysis

All diet data as analysed from droppings were averaged per brood per day for further analyses. We expressed the diet data (1) as the average relative number of barcode reads of each arthropod family in each bird species and (2) as the percentage of samples for each shorebird species where the traces of each arthropod family were found. We compared the contribution of key prey types between shorebird species using the Mann-Whitney U-test. The p-value was adjusted using the Bonferroni correction.

We estimated the date of peak arthropod abundance as the sampling date with the highest total insect biomass in the traps in relation to other sampling dates. We also estimated the dates of the peaks for two specific arthropod families, Tipulidae and Chironomidae, which turned out to be the main prey for all four shorebird species studied. We calculated the trophic mismatch as the difference between the median hatch date for each shorebird species and the arthropod peak, using both the total arthropod peak as well as the peak for the arthropod family which was the top contributor to the diet for each shorebird species.

## Results

### Arthropods sampled in the traps

The core of the arthropod community as found in pitfall traps consisted of Diptera (11 families), Hymenoptera (4 families), Coleoptera (3 families) and Araneae (families of this group were combined) (table 1). The families which contributed to over 5% of total biomass were the same in both 2018 and 2019: Muscidae (39% and 25%), Tipulidae (21% and 23%), different families of Araneae (16% and 20%), Empididae (7% and 11%) and Mycetophilidae (6% and 8%). Chironomidae, shown to be an important prey item for shorebird chicks (see the next section), contributed <1% in 2018 and 2% in 2019.

**Table 1.**
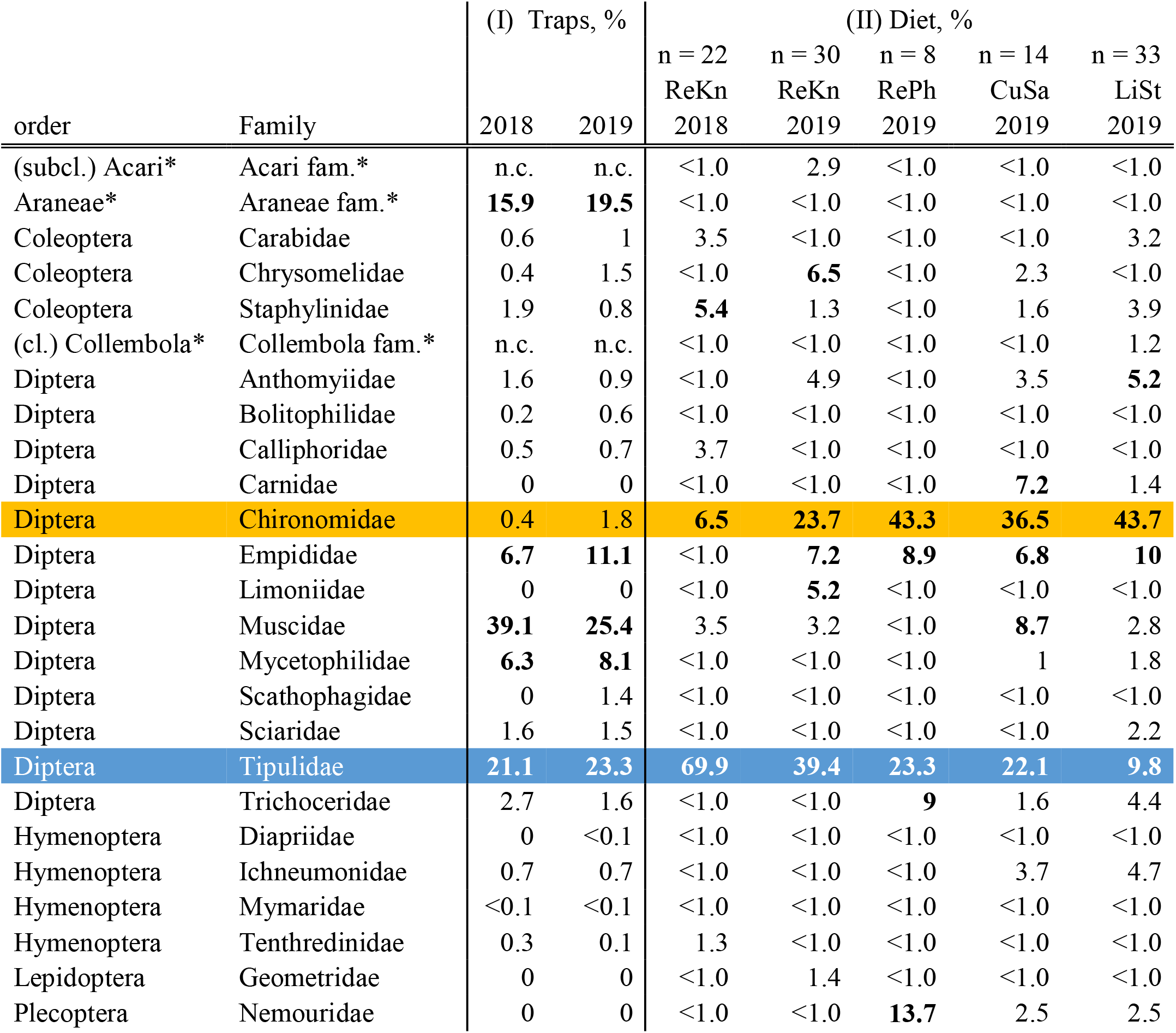
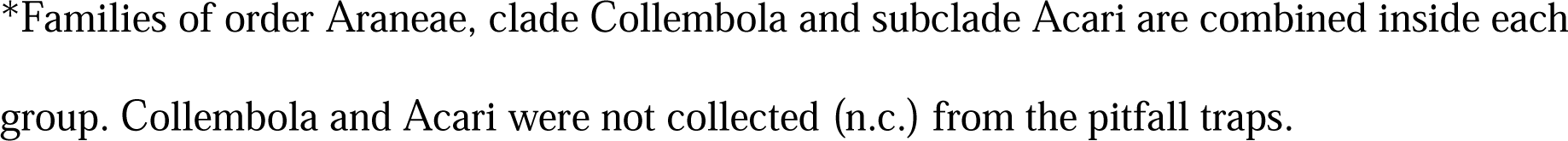
(I) The relative contribution (%) of different arthropod families to the total biomass found in the pitfall traps. (II) Average (mean) relative number of barcode reads in each shorebird species for each arthropod family. Bird species are abbreviated as ReKn – red knot, RePh – red phalarope, CuSa – curlew sandpiper and LiSt – little stint. Chironomidae and Tipulidae are marked in orange and blue, respectively. Arthropod families that contribute more than 5% are marked in bold. Arthropod families, which are presented by <0.1% in the diet in all for species and are absent from the pitfall traps are not shown.

| order | Family | (I) Traps, % |  | (II) Diet, % |  |  |  |  |
| --- | --- | --- | --- | --- | --- | --- | --- | --- |
|  |  | 2018 | 2019 | n = 22 | n = 30 | n = 8 | n = 14 | n = 33 |
|  |  |  |  | ReKn | ReKn | RePh | CuSa | LiSt |
|  |  | 2018 | 2019 | 2018 | 2019 | 2019 | 2019 | 2019 |
| (subcl.) Acari* | Acari fam.* | n.c. | n.c. | <1.0 | 2.9 | <1.0 | <1.0 | <1.0 |
| Araneae* | Araneae fam.* | **15.9** | **19.5** | <1.0 | <1.0 | <1.0 | <1.0 | <1.0 |
| Coleoptera | Carabidae | 0.6 | 1 | 3.5 | <1.0 | <1.0 | <1.0 | 3.2 |
| Coleoptera | Chrysomelidae | 0.4 | 1.5 | <1.0 | **6.5** | <1.0 | 2.3 | <1.0 |
| Coleoptera | Staphylinidae | 1.9 | 0.8 | **5.4** | 1.3 | <1.0 | 1.6 | 3.9 |
| (cl.) Collembola* | Collembola fam.* | n.c. | n.c. | <1.0 | <1.0 | <1.0 | <1.0 | 1.2 |
| Diptera | Anthomyiidae | 1.6 | 0.9 | <1.0 | 4.9 | <1.0 | 3.5 | **5.2** |
| Diptera | Bolitophilidae | 0.2 | 0.6 | <1.0 | <1.0 | <1.0 | <1.0 | <1.0 |
| Diptera | Calliphoridae | 0.5 | 0.7 | 3.7 | <1.0 | <1.0 | <1.0 | <1.0 |
| Diptera | Carnidae | 0 | 0 | <1.0 | <1.0 | <1.0 | **7.2** | 1.4 |
| **Diptera** | **Chironomidae** | **0.4** | **1.8** | **6.5** | **23.7** | **43.3** | **36.5** | **43.7** |
| Diptera | Empididae | **6.7** | **11.1** | <1.0 | **7.2** | **8.9** | **6.8** | **10** |
| Diptera | Limoniidae | 0 | 0 | <1.0 | **5.2** | <1.0 | <1.0 | <1.0 |
| Diptera | Muscidae | **39.1** | **25.4** | 3.5 | 3.2 | <1.0 | **8.7** | 2.8 |
| Diptera | Mycetophilidae | **6.3** | **8.1** | <1.0 | <1.0 | <1.0 | 1 | 1.8 |
| Diptera | Scathophagidae | 0 | 1.4 | <1.0 | <1.0 | <1.0 | <1.0 | <1.0 |
| Diptera | Sciaridae | 1.6 | 1.5 | <1.0 | <1.0 | <1.0 | <1.0 | 2.2 |
| **Diptera** | **Tipulidae** | **21.1** | **23.3** | **69.9** | **39.4** | **23.3** | **22.1** | **9.8** |
| Diptera | Trichoceridae | 2.7 | 1.6 | <1.0 | <1.0 | **9** | 1.6 | 4.4 |
| Hymenoptera | Diapriidae | 0 | <0.1 | <1.0 | <1.0 | <1.0 | <1.0 | <1.0 |
| Hymenoptera | Ichneumonidae | 0.7 | 0.7 | <1.0 | <1.0 | <1.0 | 3.7 | 4.7 |
| Hymenoptera | Mymaridae | <0.1 | <0.1 | <1.0 | <1.0 | <1.0 | <1.0 | <1.0 |
| Hymenoptera | Tenthredinidae | 0.3 | 0.1 | 1.3 | <1.0 | <1.0 | <1.0 | <1.0 |
| Lepidoptera | Geometridae | 0 | 0 | <1.0 | 1.4 | <1.0 | <1.0 | <1.0 |
| Plecoptera | Nemouridae | 0 | 0 | <1.0 | <1.0 | **13.7** | 2.5 | 2.5 |

\*Families of order Araneae, clade Collembola and subclade Acari are combined inside each
group. Collembola and Acari were not collected (n.c.) from the pitfall traps.

### Chick diet composition

Using metabarcoding to quantify the relative abundance of arthropod families in the diet, we found that two arthropod families, Tipulidae and Chironomidae, on average contributed >50% to a chick’s diet (table 1, fig. 1A). These two families occurred in different proportions in the diet of the studied shorebird species, with red knots chicks preying mainly on Tipulidae (70% in 2018 and 39% in 2019) and less on Chironomidae (7% in 2018 and 24% in 2019), whereas chicks of the other three shorebird species were mainly preying on Chironomidae (43% for red phalarope, 37% for curlew sandpiper and 44% for little stint, table 1).

**Fig. 1.**
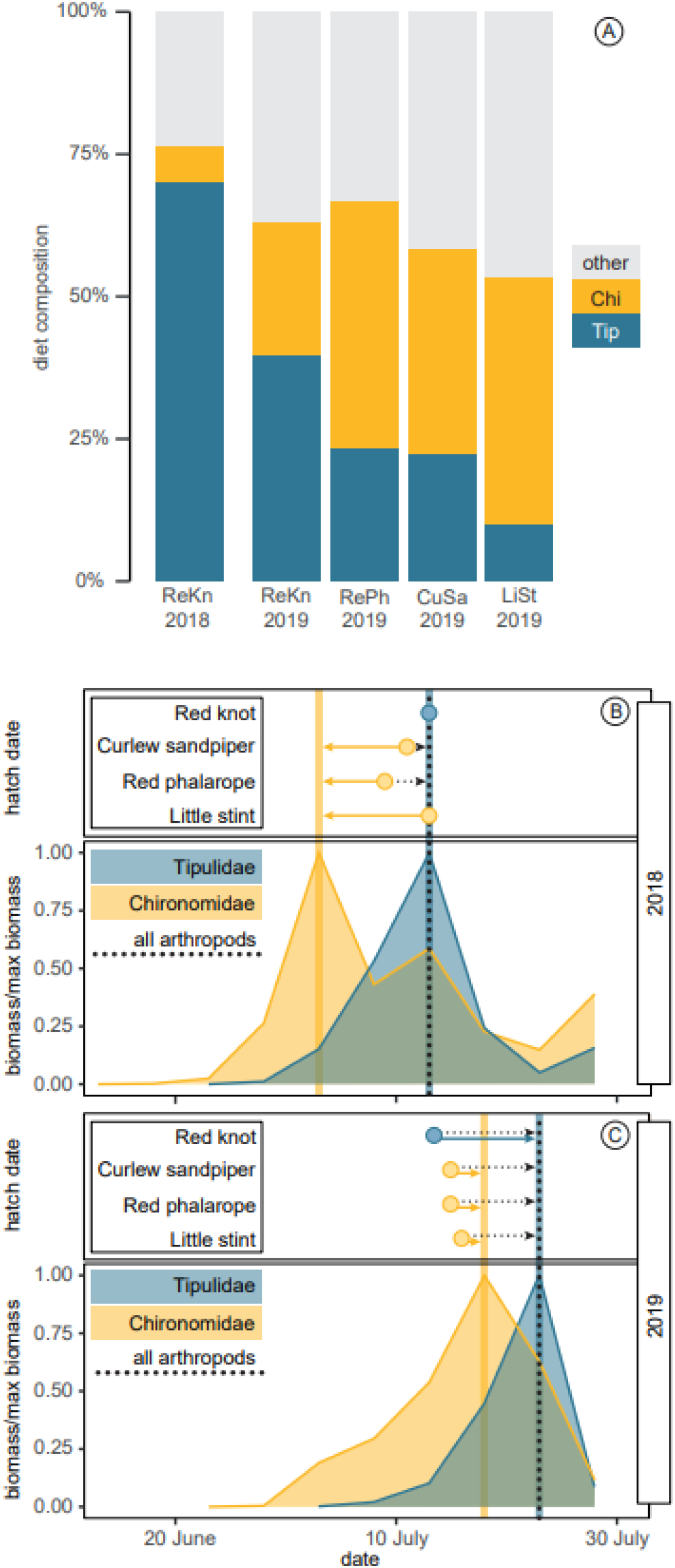
(A) The average relative number of barcode reads of Chironomidae (Chi; orange), Tipulidae (Tip; blue) and other arthropod families (other; grey) in the faeces samples of chicks of different shorebird species. ReKn – red knot, RePh – red phalarope, CuSa – curlew sandpiper, LiSt – little stint. The number below the abbreviation indicates the year. For details see table 3. (B, C) Comparison of the median hatch dates of shorebirds with the date of the total arthropod biomass peak (vertical black dotted line) and the date of the peak of arthropod prey predominating in the chick diet, including Tipulidae (vertical blue line) and Chironomidae (vertical orange line). The length and direction of the arrows indicate the degree and direction of mismatch. E.g., in 2018 curlew sandpipers hatched after the peak of Chironomidae, but before the total arthropod biomass peak, but in 2019 they hatched before both the Chironomidae and overall arthropod peaks.

These data match our results for the presence/absence of arthropod families in diet samples, with Tipulidae present in 95% of red knot samples in 2018 and 100% in 2019 (and Chironomidae in 81% and 90% of samples), whereas traces of Chironomidae were found in 100% of the samples of the other three shorebird species (S4).

Across all four species we found a negative correlation between the abundance of Tipulidae and Chironomidae, suggesting that the low abundance of Tipulidae in the diet is partly compensated by the high abundance of Chironomidae (Spearman correlation test, r = -0.59, p<0.0001). In 2019 the contribution of Tipulidae (U-test, p=0.0001, adjusted p-value=0.0083), as well as the contribution of Chironomidae (U-test, p=0.0022, adjusted p-value=0.0083) to the diet of red knots differed from those in little stints. We found no differences in the contribution of these families between other shorebirds species.

### Phenology and mismatches

The peak of the total arthropod biomass occurred on 13 July in 2018 and on 23 July in 2019. The Tipulidae peak coincided with the total arthropod peak in both years, while the Chironomidae peak occurred 10 and 5 days earlier, respectively (fig. 1B, 1C). The chicks of all shorebird species hatched around the same date, with median hatch dates being slightly earlier in 2018: red knots 13 (12 – 17 interquartile range) July, red phalarope 9 (9 – 11.5) July, curlew sandpiper 11 (7.25 – 12) July and little stint 13 (11 – 17) July; and slightly later in 2019: red knots 13.5 (7.5 – 15.75) July, red phalarope 15 (12 – 17) July, curlew sandpiper 15 (13.25 – 17.75) July and little stint 16 (14 – 19) July (fig. 1B, 1C).

Relative to the overall peak of arthropod abundance, chicks generally hatched at the moment of peak abundance in 2018 (red knots and little stints at the peak, red phalaropes and curlew sandpipers 4 and 2 days before the peak) and 1 week before the peak in 2019 (red knots 10.5 days before, red phalaropes and curlew sandpipers 8 days before and little stints 7 days before). For red knot chicks, measuring the mismatch relative to the peak of Tipulidae (the main prey) did not change the degree of mismatch. On the other hand, little stints, red phalaropes and curlew sandpiper chicks hatched after the peak of Chironomidae abundance in 2018 (red phalarope 6 days, curlew sandpiper 8 days and little stint 10 days after) and only a few days before the peak in 2019 (red phalarope 3 days, curlew sandpiper 3 days and little stint 2 days before, fig. 1B).

## Discussion

We found that the diet composition of the chicks of shorebird species varies, with different species of shorebirds selecting different arthropod prey. For shorebird species foraging mostly on Chironomidae, which emerge earlier than most other arthropods, this resulted in a potential underestimation of the mismatch between chick hatch and peak abundance of their arthropod prey by more than a week.

### Shorebird chick diet

Among all the arthropod groups that contributed the most to the total biomass during the growth period of shorebird chicks, Tipulidae and Chironomidae were the most important prey items for the chicks. In contrast, some groups, e.g., spiders (Araneae) and house flies (Muscidae), which were abundant in pitfall traps, were rarely found in the diet of shorebird chicks. There may be several explanations for these patterns, including (1) the chicks’ preferences for some arthropods, if they are easier prey than others, (2) a potential bias in sampling with pitfall traps by catching more arthropods of certain species, (3) the selectivity of the applied molecular-genetic method for certain families (Deagle et al. 2014).

As suggested by other studies, the family Tipulidae that includes large crane fly species, plays a key role in the food supply of Arctic shorebirds (Rakhimberdiev 2007). In contrast to, e.g., Muscidae, the Tipulidae seem to be the poorer flyers, and a large part of the individuals in our samples is represented by wingless, ground-dwelling morphs. Their large size makes Tipulidae a clearly visible and profitable prey which would explain a high preference by shorebird chicks. Chironomidae have also been considered to be an important and highly abundant prey for shorebird chicks (Gerik 2018; Drury 1961), especially so in Arctic habitats (MacLean, Jr. and Pitelka 1971; Hodkinson et al. 1996). The relatively low abundance found in our study site is likely explained by the use of pitfall traps, since studies using other trapping methods such as Malaise traps or boards covered with sticky resin generally found much higher abundances of Chironomids (MacLean, Jr. and Pitelka 1971).

Although Tipulidae and Chironomidae are the key prey items for the chicks of all species of shorebirds and together make up 54% -76% of the chick diet, their fraction in the diet varies, and the absence of one family is compensated by the presence of the other. This variation may in part be explained by morphometric differences between bird species and their chicks. Compared with little stints, red knot chicks are up to 4 times as large at the age of 10 days (Tjørve et al. 2007; Lameris et al. 2022) which may explain why they consumed relatively more of the larger Tipulidae and less of the smaller Chironomidae. Larger Tipulidae may be more profitable and lead to higher intake rates (Stephens and Krebs 1986), but only for shorebird chicks with bills and digestive system large enough to handle such large prey. On top of this, the variation in the relative amount of Chironomidae and Tipulidae in the diet appears to be also explained by annual differences in availability, as red knots consumed more Chironomidae in 2019, when some chicks hatched before the peak in Tipulidae abundance. Unfortunately, we were unable to sample droppings from the other shorebird species in 2018 and therefore do not know whether Chironomidae were also the preferred prey in that year, or whether Tipulidae would also have been more common in the diet.

### Implications for trophic mismatches

Our results show that arthropod species that form the basis of the diet differ in chicks of the different shorebird species, and this has important implications for whether their growth period matches the peak in abundance of this prey. As the peak in abundance of Tipulidae coincides with the general peak in arthropod abundance, this does not affect the interpretation of a mismatch for red knot chicks. On the other hand, the peak in abundance of Chironomids falls 5 – 10 days before the main arthropod peak, and as such the mismatch as interpreted for red phalarope, curlew sandpiper and little stint chicks differed (larger mismatch in 2018, hatching shortly before the food peak in 2019) than when considering the main arthropod peak. Our results suggest that, when detailed information on the chick diet is lacking, it is difficult to correctly interpret the degree of trophic mismatch (Samplonius et al. 2016) and therefore to analyse whether or not trophic mismatches have fitness repercussions. As no information on the diet is available for most shorebird chicks, the degree of trophic mismatches may often be over- or underestimated.

