## Supplementary material for "Disentangling the diet composition of Arctic shorebirds’ chicks provides a new perspective on trophic mismatches": S1

*Barcoding method description*

DNA extraction and PCR amplification
Genomic DNA was extracted from the 180 fecal samples using a PureLink™ Microbiome DNA Purification Kit (Invitrogen™, UK) following the manufacturer protocol with small modifications. Entire fecal samples (average wet weight 240 (±180) mg) were transferred to a pre-weighted UV-cleaned tube using a metal scoop. For big samples (heavier than 1 g) a subsampling approach was used to prevent PCR inhibition by uric acids. To avoid contamination, after each sample gloves were changed, a desk was cleaned with RNase away and a metal scoop was cleaned and flamed. To remove all the ethanol from the sample, it was dried at 55°C for ca. 30 min. After drying, the samples were weighted (average dry weight 22 (±19) mg) transferred to PureLink bead tubes using autoclaved and UV-cleaned toothpicks. A small amount of 0.1 mm Zircona/Silica beads was additionally added to each bead-tube with the sample. The samples were homogenized at 5.00 m/s, 2 cycles for 10 s with a 30 s break in 700 µl of lysis buffer provided with the kit, using a Bead Ruptor Elite homogenizer (Omni International, USA). For each DNA-extraction set, one negative control was applied. The primer pairs, LCO1490_5T (5′-GGTCTACAAATCATAAAGATATTGG-3′) and
HCO1777_15T (5′-ACTTATATTATTTATACGAGGGAA-3′) flanking a 304-bp fragment of the standard 710-bp cytochrome C oxidase (COI) gene barcoding region were used. These primers were based on general invertebrate cytochrome oxidase I primers LCO1490 (Folmer, Black, Hoeh, Lutz, & Vrijenhoek, 1994) and HCO1777 (Brown, Jarman, & Symondson, 2012) and modified by Verkuil et al., 2022 to reduce bird host reads to 0.03%, and amplify Arachnida DNA without significant changing the recovery of other arthropod taxa. Primers that amplify short fragments such as this are needed to ensure PCR success from semi-digested samples (King, Read, Traugott, & Symondson, 2008; Symondson, 2002).

All PCRs were performed in a reaction mix containing 2 µl of DNA template, 2.9 µl of each primer (10 μM), 14.5 µl of AccuStart™ II PCR Tough Mix (Quantabio, USA), fluorescent dye Eva Green 0.7 µl and made up to 25 µl with water. Triplicate reactions were performed for each sample. Each sample received a separate primer pair with two unique 12-base barcodes on forward and reverse primer (Integrated DNA Technologies, USA).
Each reaction set included 6 standard samples and 2 negative PCR-controls.
The thermocycling profile was according to Verkuil et al., 2022 and included: initial denaturation at 94°C for 2 min 30 s, followed by 35 cycles of denaturation at 94°C for 30 s, annealing at 48°C for 30 s, elongation at 72°C for 45 s; and a final elongation at 72°C for 10 min. The Qiagen Rotor-gene qPCR thermocycler (Qiagen, USA) was used for all the amplifications.

All PCR products were checked on 2% agarose gel. Subsequently, the triplicates were pooled and quantitative gels were run. On these gels three quantification standards (20 ng/µl, 10 ng/µl and 1 ng/µl) were run along with the samples allowing calculation of PCR product concentration based on pixel densities.

To obtain an equal distribution of sequencing reads among samples, pooling all the samples in equimolar amounts was performed. The final pooled sample was gel-extracted with QIAquick Gel Extraction Kit (Qiagen, USA) and further purified and concentrated using a QIAquick PCR Purification Kit. The final concentration was measured on a Qubit machine (Invitrogen™).
Illumina Mi-Seq PE300 sequencing was performed at USEQ (Utrecht Sequencing facility, The Netherlands) using TrueSeq library preparation and the V3 kit.

*Potential bias of the applied molecular-genetic method*

In order to minimize bias caused by the molecular methods, we applied the same protocols as used by Verkuil et al. 2022 who showed a good correlation between relative read abundance of insect prey and camera footage. In addition, we tested whether the primers in combination with the low annealing temperature applied were able to amplify all arthropod families that were caught with the traps. As a result all trapped arthropod families produced PCR products. Nevertheless, in-silico analyses of the number of mismatches with the reverse primer indicates potential bias (>5 mismatches) for some Linyphiidae genera (Hybauchenidium, Collinsia, Incestophantes and Pityohyphantes) and many Lycosidae.

**Acknowlegments**

We thank Harry Witte for running the bioinformatic analyses and Hans Malschaert for being a Linux helpdesk.
