## Supplementary material for "Disentangling the diet composition of Arctic shorebirds’ chicks provides a new perspective on trophic mismatches": S4

Table. Presence of the arthropod families in the faeces samples of shorebird chicks, as indicated by metabarcoding method. Bird species are abbreviated as ReKn – red knot, RePh – red phalarope, CuSa – curlew sandpiper and LiSt – little stint. The number above abbreviation is the year when samples were collected. The data are given in % of the total number of samples of a shorebird species, in which the traces of arthropod family were found. Chironomidae and Tipulidae are marked with the same colours as in figures 1. Arthropod families which were found in more than 50% of samples are marked in bold. *Families of Acari, Araneae and Collembola are combined inside each group. **The column order incudes also subclade Acari and clade Collembola, indicated by (subcl.) and (cl.).

|  |  | 2018 | 2019 | 2019 | 2019 | 2019 |
| --- | --- | --- | --- | --- | --- | --- |
|  |  | ReKn | ReKn | RePh | CuSa | LiSt |
| order | family | n = 22 | n = 30 | n = 8 | n = 14 | n = 33 |
| (subcl.) Acari** | Acari families* | 33 | 33 | 13 | 29 | 21 |
| Diptera | Agromyzidae | 10 | 0 | 0 | 14 | 3 |
| Diptera | Anthomyiidae | 10 | **77** | **75** | **100** | **88** |
| Trichoptera | Apataniidae | 0 | 3 | 0 | 0 | 0 |
| Araneae | Araneae families* | 48 | 30 | 38 | **50** | **58** |
| Hymenoptera | Braconidae | 0 | 0 | 0 | 7 | 3 |
| Diptera | Calliphoridae | 5 | 0 | 0 | 0 | 3 |
| Coleoptera | Carabidae | 10 | 7 | 0 | 7 | 6 |
| Diptera | Carnidae | 0 | 10 | 0 | 36 | 18 |
| Diptera | Cecidomyiidae | 0 | 7 | 0 | 14 | 18 |
| Diptera | Chironomidae | **81** | **90** | **100** | **100** | **100** |
| Coleoptera | Chrysomelidae | 48 | **53** | 25 | 36 | 21 |
| Coleoptera | Coccinellidae | 5 | 0 | 0 | 0 | 0 |
| (cl.) Collembola** | Collembola families* | **52** | 33 | 25 | 29 | 48 |
| Diptera | Culicidae | 0 | 0 | 0 | 14 | 0 |
| Diptera | Dolichopodidae | 0 | 3 | 0 | 0 | 3 |
| Coleoptera | Dytiscidae | 0 | 0 | 25 | 0 | 6 |
| Diptera | Empididae | 48 | **80** | **88** | **100** | **85** |
| Diptera | Ephydridae | 5 | 10 | 13 | 14 | 0 |
| Lepidoptera | Geometridae | 0 | 7 | 0 | 14 | 15 |
| Coleoptera | Haliplidae | 0 | 0 | 0 | 0 | 9 |
| Hymenoptera | Ichneumonidae | 29 | 20 | 25 | 36 | **55** |
| Diptera | Limoniidae | 19 | 27 | 0 | 7 | 3 |
| Diptera | Milichiidae | 5 | 17 | 0 | 0 | 3 |
| Diptera | Muscidae | **52** | **73** | **88** | **93** | **97** |
| Diptera | Mycetophilidae | 38 | **57** | **75** | **93** | **91** |
| Plecoptera | Nemouridae | 5 | 0 | 38 | 14 | 12 |
| Lepidoptera | Noctuidae | 14 | 17 | 25 | 21 | 6 |
| Diptera | Phoridae | 0 | 7 | 0 | 14 | 3 |
| Diptera | Piophilidae | 0 | 0 | 0 | 0 | 3 |
| Lepidoptera | Plutellidae | 0 | 0 | 0 | 0 | 3 |
| Lepidoptera | Pterophoridae | 5 | 0 | 0 | 0 | 0 |
| Lepidoptera | Pyralidae | 0 | 0 | 13 | 0 | 0 |
| Diptera | Rhagionidae | 0 | 0 | 0 | 7 | 0 |
| Diptera | Sarcophagidae | 0 | 23 | 25 | **50** | 21 |
| Diptera | Scathophagidae | 14 | 27 | **63** | 43 | **58** |
| Diptera | Sciaridae | 24 | 20 | 25 | **50** | 48 |
| Diptera | Simuliidae | 10 | 3 | 0 | 7 | 0 |
| Diptera | Sphaeroceridae | 14 | 3 | 0 | 7 | 0 |
| Coleoptera | Staphylinidae | 29 | 43 | 38 | 36 | 36 |
| Diptera | Syrphidae | 0 | 17 | 25 | 36 | 39 |
| Diptera | Tachinidae | 0 | 0 | 13 | 7 | 3 |
| Hymenoptera | Tenthredinidae | 10 | 33 | 13 | 21 | 24 |
| Diptera | Tipulidae | **95** | **100** | **75** | **79** | **61** |
| Lepidoptera | Tortricidae | 19 | 33 | 13 | 14 | 0 |
| Diptera | Trichoceridae | 14 | 10 | **63** | **57** | 48 |
